# “Targeting LARP1 Enhances Carboplatin Sensitivity and Suppresses Tumor Growth in Endometrial Cancer”

**DOI:** 10.64898/2026.03.22.713473

**Authors:** Abdelrahman M. Elsayed, Marwa W. Eldegwy, Salama A. Salama

## Abstract

La-related protein 1 (LARP1) is an RNA-binding protein that post-transcriptionally regulates mRNA with potential oncogenic role in multiple cancers; however, its function in endometrial cancer remains unknown. An analysis of the TCGA endometrial cancer cohort showed that overexpression of LARP1 is associated with shorter overall survival (OS) and progression-free interval (PFI) as indicated by Kaplan-Meier analysis. Functional *in vitro* studies revealed that LARP1 knockdown by two different siRNAs markedly suppressed cell viability and triggered apoptosis, as confirmed by increased protein levels of cleaved PARP1 and cleaved caspase-3. Mechanistically, LARP1 knockdown remarkably reduced E2F1 protein levels as confirmed by immunofluorescence and Western blotting. Clinically, co-overexpression of LARP1 and E2F1 further decreased OS and PFI, suggesting a co-operative oncogenic axis. Importantly, LARP1 knockdown enhanced the sensitivity of ISHI and HEC-1A endometrial cancer cell lines to carboplatin treatment. These findings suggest that LARP1 promotes endometrial cancer survival and resistance to chemotherapy, at least in part, through the regulation of E2F1 and suppression of apoptosis. Targeting LARP1 could represent a promising therapeutic strategy to suppress tumor growth and enhance sensitivity to platinum-based chemotherapy.

## 1. Introduction

Endometrial cancer (EC) is one of the most prevalent gynecologic malignancies worldwide [1]. In the United States, endometrial cancer remains the most common invasive gynecologic malignancy and a growing public health concern. In 2026, an estimated 68,270 new cases and 14,450 deaths are projected. Approximately 68% of cases are diagnosed at an early stage, largely due to symptomatic presentation. Although overall incidence rates have stabilized among White women, they continue to increase by 1.8–2.6% annually among women of other racial and ethnic groups. Mortality rates have also risen over the past decade, increasing by approximately 1.5–1.6% per year [2]. Advanced and recurrent disease presents substantial therapeutic challenges, as response rates to conventional treatment modalities remain limited. Systemic chemotherapy is generally indicated for individuals with disseminated primary disease or extra pelvic recurrence. Carboplatin–paclitaxel regimen is currently the standard first-line therapy for the treatment of advanced or recurrent endometrial cancer [3]. However, the development of resistance is a major challenge that can result in disease recurrence and poor clinical outcome [4, 5]. Multiple molecular mechanisms have been reported to mediate platinum resistance. These include decreased intracellular drug accumulation through altered expression of membrane transporters and enhanced detoxification, and improved DNA damage repair machinery [4]. In addition, cancer cells may evade the cytotoxic action of platinum agents by upregulating pro-survival signaling pathways such as PI3K/AKT and MAPK or downregulating apoptotic pathways [6]. Furthermore, emerging evidence indicates that aberrant expression of long-noncoding RNAs and RNA-binding proteins (RBPs) can promote resistance to platinum-based chemotherapy [7-10]. However, the role of La-related protein 1 (LARP1), an RNA-binding protein, in endometrial cancer remains unclear.

Dysregulation of RBPs has been shown to promote tumor growth, metastasis, apoptosis evasion, and resistance to therapy in endometrial cancer [11]. RBPs play a critical role in regulating gene expression at the post-transcriptional level; they bind to different forms of RNA,forming ribonucleoprotein complexes that regulate RNA stability, decay, splicing, localization, and protein translation [12]. LARP1 is a highly conserved RBP that was first identified in *Drosophila melanogaster* and has been shown to interact with poly(A)-binding protein and orchestrate embryonic development and fertility [13]. Subsequent studies revealed that LARP1 regulates the stability and translation of 5′-terminal oligopyrimidine (TOP) mRNAs, a class of mRNAs required for production of ribosomal proteins and translational factors [14, 15]. Previous studies demonstrated that LARP1 protein is overexpressed in multiple cancers, including hepatocellular carcinoma, and lung, ovarian, and cervical cancer, where it serves as an independent predictor of poor prognosis [15-17]. However, the role of LARP1 in endometrial cancer remains unknown. Here, we report that LARP1 is overexpressed in endometrial cancer patients and associated with shorter overall survival (OS) and progression-free interval (PFI) and that targeting LARP1 inhibits survival, induces apoptosis, and sensitizes the response to carboplatin *in vitro*.

## 2. Materials and methods

### 2.1. TCGA Endometrial Cancer Cohort Data Analysis

Analysis of TCGA data was performed using R software (version: 2025.09.0). Statistical significance was defined as p < 0.05. Clinical information for patients with endometrial cancer and RNA-sequencing data (FPKM-UQ normalized) were retrieved from the Genomic Data Commons Data Portal. After excluding samples lacking either clinical information or gene expression data, 454 primary endometrial tumor samples were included in the final analysis.

For survival analyses, patients were stratified into high- and low-expression groups based on the median expression value of LARP1 or E2F1. Survival distributions were estimated using the Kaplan–Meier method, and differences between groups were evaluated using the log-rank test. The Spearman rank correlation test was used to evaluate the association between the mRNA expression levels of LARP1 and E2F1. To assess the prognostic significance of gene expression and clinical variables, univariate Cox proportional hazards regression models were applied for overall survival (OS) and progression-free interval (PFI). Multivariate Cox proportional hazards model was applied for OS and PFI and included age at diagnosis, disease stage, as well as LARP1 or E2F1.

### 2.2. Cell culture and siRNA transfection

Human endometrial cancer cell lines HEC-1A and Ishikawa (ISHI) were obtained from ATCC (American Type Culture Collection, Manassas, VA, USA). HEC-1A cells were cultured in McCoy’s 5A supplemented with 10% fetal bovine serum (FBS) and 1% penicillin/streptomycin (P/S), whereas ISHI cells were cultured in MEM supplemented with 10% FBS and 1% P/S. All cell lines were maintained in 5% CO_2_ and 95% air at 37 °C. All cell lines were screened for mycoplasma using the MycoAlert mycoplasma detection kit (Lonza, Basel, Switzerland). All experiments were conducted with cell cultures at 60–80% confluency. All siRNA transfections were performed using lipofectamine 3000 (Thermo Fisher Scientific, Waltham, MA, USA) according to the manufacturer’s protocol. Two independent sequences of LARP1 siRNA and a negative control siRNA were used. The sequences of siRNAs used are shown in **Table 1**. Different concentrations of LARP1 siRNA (25, 50, 75, or 100 nM) were mixed with optimum and lipofectamine 3000, incubated at room temperature for 15 min, and added dropwise to HEC-1A or ISHI cells plated 24 h earlier. Cells were then harvested at 72 – 96 h following transfection unless otherwise specified.

**Table 1:**
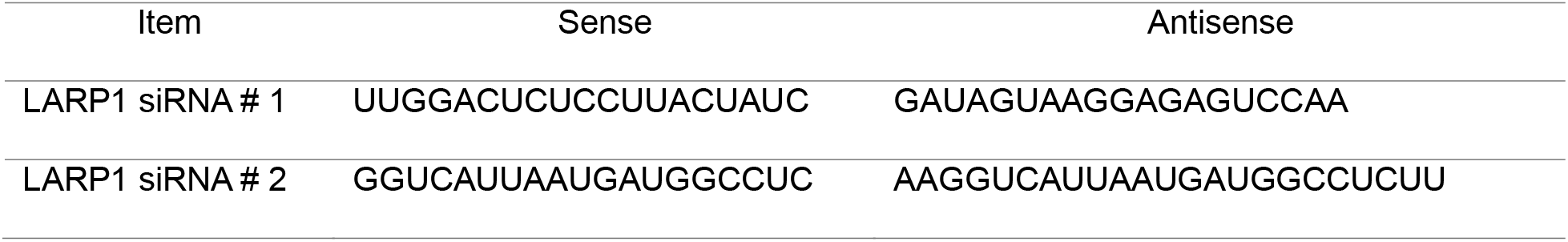
List of siRNA sequences.

### 2.3. MTS cell viability assay

Cell viability was assayed using the MTS reagent. Briefly, HEC-1C and ISHI (2× 10^3^/well) cells were seeded in 96-well culture plates. Cells were treated with escalating concentrations of LARP1 siRNA (25, 50, 75, 100 nM) and allowed to grow at regular culture conditions (5% CO_2_, 37 °C, and 95% air). At 96 h following transfection, 20 µL of MTS reagent (Promega Corporation, Madison, WI, USA) was added to each well and the absorbance was recorded at 490 nm by a spectrophotometer (Agilent Technologies, Santa Clara, CA, USA). For carboplatin (Pfizer, New York, NY, USA) experiment, cells were plated in 96-well plate and treated with escalating concentrations of carboplatin (12.5 – 400 µM). At the end of treatment schedule, cells were mixed with 20 µL of MTS reagent and the absorbance was recorded at 490 nm by a spectrophotometer (Agilent Technologies, Santa Clara, CA, USA).

### 2.4. Cell extracts and Western blot analysis

A total cell pellet was lysed in standard radioimmunoprecipitation assay (RIPA) buffer containing protease and phosphatase inhibitor cocktail (Roche, Basel, Switzerland). To purify the lysate, pellet-containing RIPA buffer was centrifuged at 14,000rpm for 20 min at 4 °C, and supernatants were collected. The protein concentration was determined using the Pierce bicinchoninic acid (BCA) protein assay kit according to the manufacturer’s protocol (Thermo Fisher Scientific, Waltham, MA, USA). Appropriate protein concentrations (40–80 µg) were mixed with 4X Laemmli loading dye, heated at 100 °C for 5 min, loaded into sodium dodecyl sulfate polyacrylamide gel (4%–15%), and transferred at 100 V for 60 min, 4 °C to polyvinylidene difluoride membranes (MilliporeSigma, Burlington, MA, USA). The expression levels of selected proteins were determined using their corresponding primary antibodies followed by incubation with horseradish peroxidase–conjugated secondary antibodies **(Table 2)**. Immunoblots were developed using SuperSignal™ Western Blot Substrate Trial Pack, Atto & Pico PLUS (Thermo Fisher Scientific, Waltham, MA, USA). The signals were recorded using Azure Imager 300 (Azure Biosystems, CA, USA). β-actin was used as a loading control.

**Table 2:**
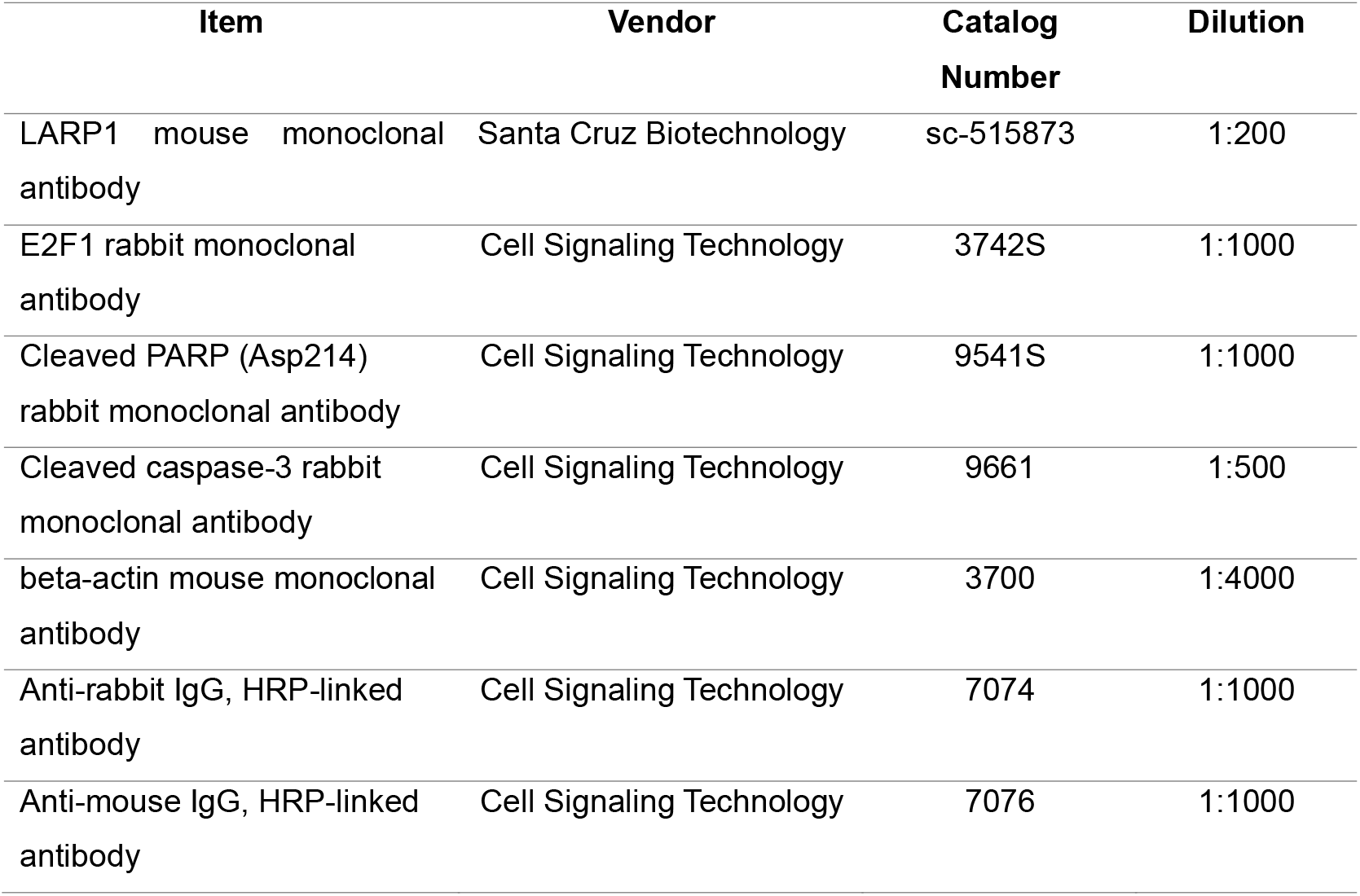
List of antibodies used in western blot.

### 2.5. Colony Formation Assay

Cells were plated onto 12-well plate (3 ×10^3^ cells/well) and transfected with control or LARP1 siRNA at 50 nM. At 24 h post-transfection, the media was replaced with fresh complete media (RPMI with 10% FBS and 1% penicillin/streptomycin). Seventy-two hours post transfection; carboplatin was added at the desired concentration (12.5 µM) and cells were allowed to grow. Old media was replaced every 3 days until the end of experiment. At the end of the treatment schedule (12 – 14 days), the colonies were washed once with phosphate-buffered saline, stained with crystal violet (0.5% w/v), and photographed. The color was extracted using 10% glacial acetic acid (MilliporeSigma, St. Louis, MO, USA) and the absorbance of the stain intensity was determined at 590 nm using Agilent Gen 5 ELISA (Agilent Technologies, Santa Clara, CA, USA). Three replicates were used to measure statistical significance between the experimental groups.

### 2.6. Immunofluorescence staining

Cells were washed with phosphate-buffered saline (PBS) (Invitrogen, Waltham, MA, USA), fixed with 4% paraformaldehyde (Sigma-Aldrich, St. Louis, MO, USA), and permeabilized with 0.05% Triton X-100 (Sigma-Aldrich). To block nonspecific binding, cells were incubated in blocking buffer consisting of 5% goat serum and 1% bovine serum albumin (BSA) in PBS for 60 min at room temperature. Cells were then incubated overnight at 4 °C with the following primary antibodies: LARP1 (Cat. No. sc-515873; 1:200 dilution; Santa Cruz Biotechnology, Dallas, TX, USA) and E2F1 (Cat. No. 3742S; 1:400 dilution; Cell Signaling Technology, Danvers, MA, USA). After washing with PBS, cells were incubated for 1 h at room temperature with secondary antibodies: goat anti-mouse IgG Alexa Fluor 488 (Cat. No. A-11001) and goat anti-rabbit IgG Alexa Fluor 568 (Cat. No. A-11011) (Invitrogen, Thermo Fisher Scientific, Waltham, MA, USA). Cells were subsequently washed with PBS and counterstained with DAPI (4′,6-diamidino-2-phenylindole) in an aqueous mounting medium (Cat. No. 00-4959-52; Invitrogen, Thermo Fisher Scientific, Waltham, MA, USA). Images were acquired using a confocal microscope (Nikon, Melville, NY, USA).

### 2.7. Statistical analysis

For comparisons involving more than two groups, one-way analysis of variance (ANOVA) followed by Tukey’s post hoc multiple comparison test was performed. All statistical analyses were conducted using GraphPad Prism (version 10.5.0; GraphPad Software, San Diego, CA, USA). Data are presented as mean ± standard deviation (SD), with individual expression values presented as dots on the bar plots. All statistical tests were two-tailed, and P ≤ 0.05 was considered statistically significant.

## 3. Results

### 3.1. LARP1 is overexpressed in endometrial cancer and associated with shorter overall and progression-free survival

Previous reports revealed that the RNA-binding protein LARP1 is a positive regulator of tumorigenesis and that overexpression of LARP1 is correlated with poor prognosis in multiple cancers [15-17]. Consistent with these findings, analysis of The Cancer Genome Atlas (TCGA) endometrial cancer (EC) dataset revealed that LARP1 overexpression is associated with shorter overall survival (OS, p = 0.0035) and reduced progression-free interval (PFI, p = 0.044; **Figure 1A, B**). Univariate Cox proportional hazards analysis further confirmed that LARP1 overexpression is correlated with increased risk of death (HR = 1.863, p = 0.004, 95% CI: 1.219 - 2.849) and disease progression (HR = 1.438, p = 0.0449, 95% CI = 1.008 - 2.052). After adjustment for age and stage in multivariate Cox regression analysis, the association between LARP1 and clinical outcome was attenuated and did not reach statistical significance in OS (HR = 1.456, p = 0.086, 95% CI: 0.947 - 2.239) and PFI (HR = 1.255, p = 0.216, 95% CI: 0.876 - 1.798). These data suggest that the impact of LARP1 on clinical outcome in EC may be partially influenced by other clinical variables. Next, we set out to knock down LARP1 in two different EC cell lines, HEC-1A and ISHI (Ishikawa). We used 4 different doses of LARP1 siRNA (25, 50, 75, and 100 nM) and showed that 75 and 100 nM produced an optimal knockdown of LARP1 protein in HEC-1A cell line and therefore were selected for further experiments **(Figure 1C)**. Immunofluorescence staining further validated the knockdown of LARP1 in HEC-1A cell line by using two different LARP1-targeting siRNAs (**Figure 1D**).

**Figure 1.**
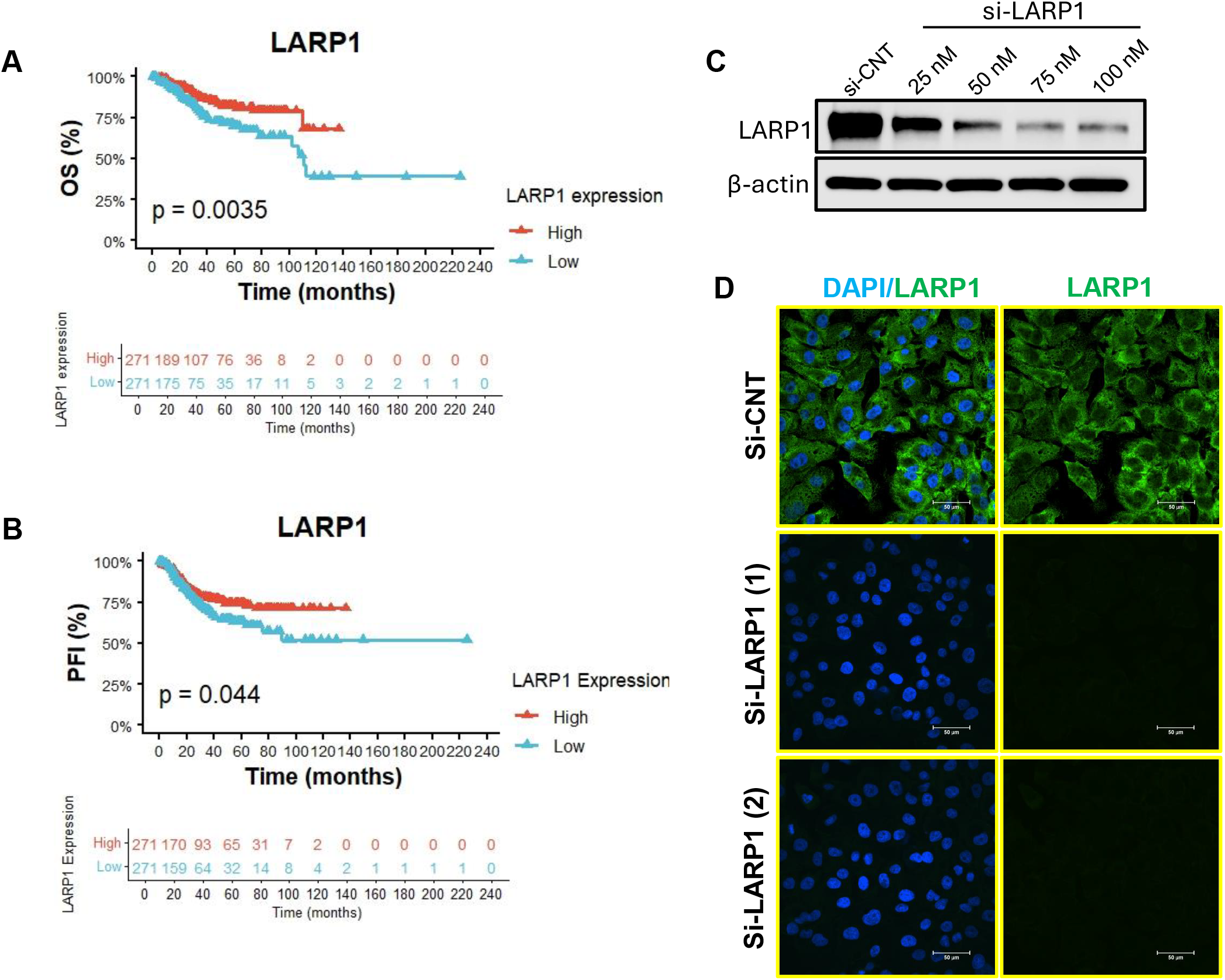
LARP1 overexpression is associated with shorter survival in TCGA endometrial cancer dataset. (A, B) Kaplan-Meier plots showing that LARP1 overexpression is associated with shorter overall survival (A) and progression-free interval (B) in endometrial cancer cohort (n=454). (C) Immunoblot analysis illustrating the protein expression of LARP1 after transfecting ISHI cell line with control or LARP1 siRNA at 25, 50, 75, and 100 nM. β-actin was used as a loading control. (D) Images derived from immunofluorescence analysis showing the protein expression of LARP1 after transfecting HEC-1A cells with control or LARP1 siRNA at 75 µM. Scale bar = 50 µM.

### 3.2. LARP1 knockdown inhibits cell viability and induces apoptosis in endometrial cancer cell lines

Given that LARP1 overexpression is associated with shorter OS and PFI in EC patients, we set out to investigate how LARP1 knockdown influences cell viability *in vitro*. We therefore induced LARP1 knockdown in two different EC cell lines and demonstrated that LARP1 knockdown significantly inhibited the viability of ISHI and HEC-1A lines compared to control siRNA-transfected cells (**Figure 2A – D**). Importantly, the suppression of cell viability was more pronounced in ISHI cell line than in HEC-1A cell line, suggesting that ISHI cells are more sensitive to LARP1 knockdown. Previous reports revealed that LARP1 suppresses apoptosis by stabilizing mRNA of key anti-apoptotic proteins [17]. We therefore performed western blot to determine the protein expression levels of cleaved PARP1 and cleaved caspase 3, well known markers of apoptosis. Our results revealed that transfecting ISHI and HEC-1A cell lines with BAP1 siRNA resulted in a remarkable elevation of cleaved PARP1 and cleaved caspase 3 protein levels compared to cells transfected with control siRNA **(Figure 2E)**. Collectively, these data suggest that LARP1 promotes EC tumorigenesis by promoting cell survival and resisting apoptosis.

**Figure 2.**
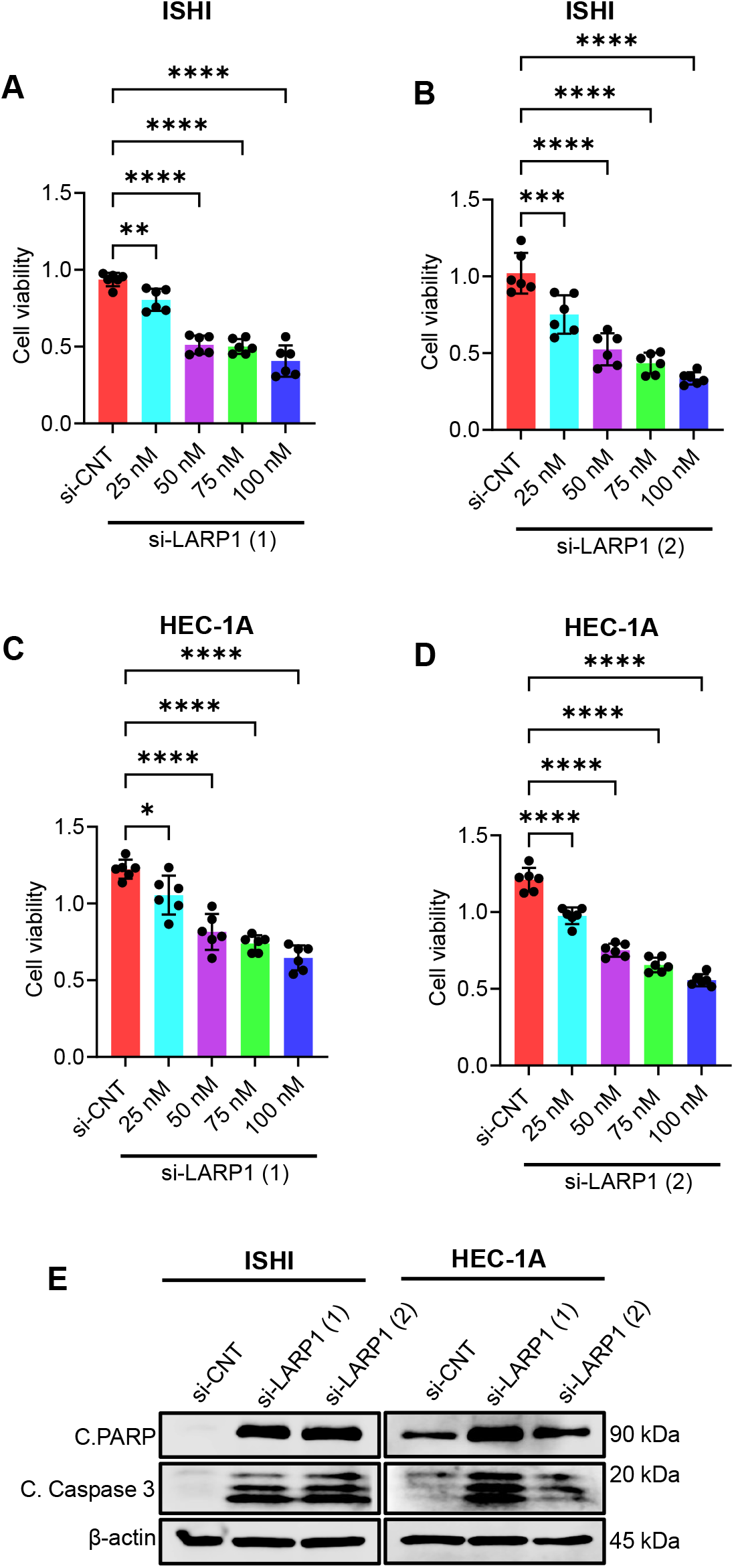
LARP1 knockdown inhibits viability and induces apoptosis of endometrial cancer cells. (A – D) Cell viability assay showing the effect of two different LARP1 siRNA (1 and 2) on the viability of ISHI (A, B) and HEC-1A (C, D) cell lines. One-way analysis of variance (ANOVA) was used to assess statistical significance among groups, followed by Tukey’s post hoc test for multiple pairwise comparisons. Data are presented as mean ± SD, with individual data points shown as dots. (E) Immunoblot analysis showing the protein expression of cleaved PARP1 and cleaved caspase 3 after transfection of ISHI and HEC-1A cells with control or LARP1 siRNAs. β-actin was used as a loading control.

### 3.3. LARP1 knockdown inhibits E2F1 protein expression

Our unpublished data derived from reverse phase protein array indicated that LARP1 knockdown inhibits the protein expression of E2F, a key regulator of cell cycle progression. We thus analyzed TCGA EC dataset and revealed that E2F1 overexpression is associated with shorter OS (p < 0.0001) and reduced PFI (p < 0.0001, **Figure 3A, B**). Univariate Cox proportional hazards analysis further confirmed that E2F1 overexpression is correlated with increased risk of death (HR = 2.366, p = 0.000152, 95% CI: 1.515 - 3.694) and disease progression (HR = 2.208, p 3.38e-05, 95% CI = 1.518 - 3.21). After adjustment for age and stage in multivariate Cox regression analysis, the association between E2F1 and clinical outcome was significant in PFI (HR = 1.675, p = 0.01, 95% CI: 1.131 - 2.481), but it was attenuated and did not reach statistical significance in OS (HR = 1.511, p = 0.0862, 95% CI: 0.943 - 2.423). Interestingly, co-overexpression of both LARP1 and E2F1 was significantly associated with a further reduction in OS (p < 0.0001) and PFI (p < 0.0001, **Figure 3C, D**). Spearman correlation analysis of EC cohort (n = 554) indicated that LARP1 is positively correlated with E2F1 (r = 0.171, p = 5.27e-05; **Figure 3E**). To investigate the association between LARP1 and E2F1 in EC cell lines, we performed western blot after LARP1 transfection in ISHI and HEC-1A cells. First, we confirmed the knockdown of LARP1 in both cell lines following the transfection with two different LARP1 siRNAs (**Figure 4A, B**). Next, we detected the protein expression level of E2F1 and demonstrated that LARP1 knockdown markedly decreased the protein expression of E2F1 in both cell lines (**Figure 4A, B**). Similar results were further reproduced using immunofluorescence analysis in HEC-1A cell line, where LARP1 knockdown reduced the protein expression level of E2F1 (**Figure 4C**). Conclusively, these data suggest that the effect of LARP1 on tumor growth and proliferation could be mediated, at least in part, by positively regulating E2F1.

**Figure 3.**
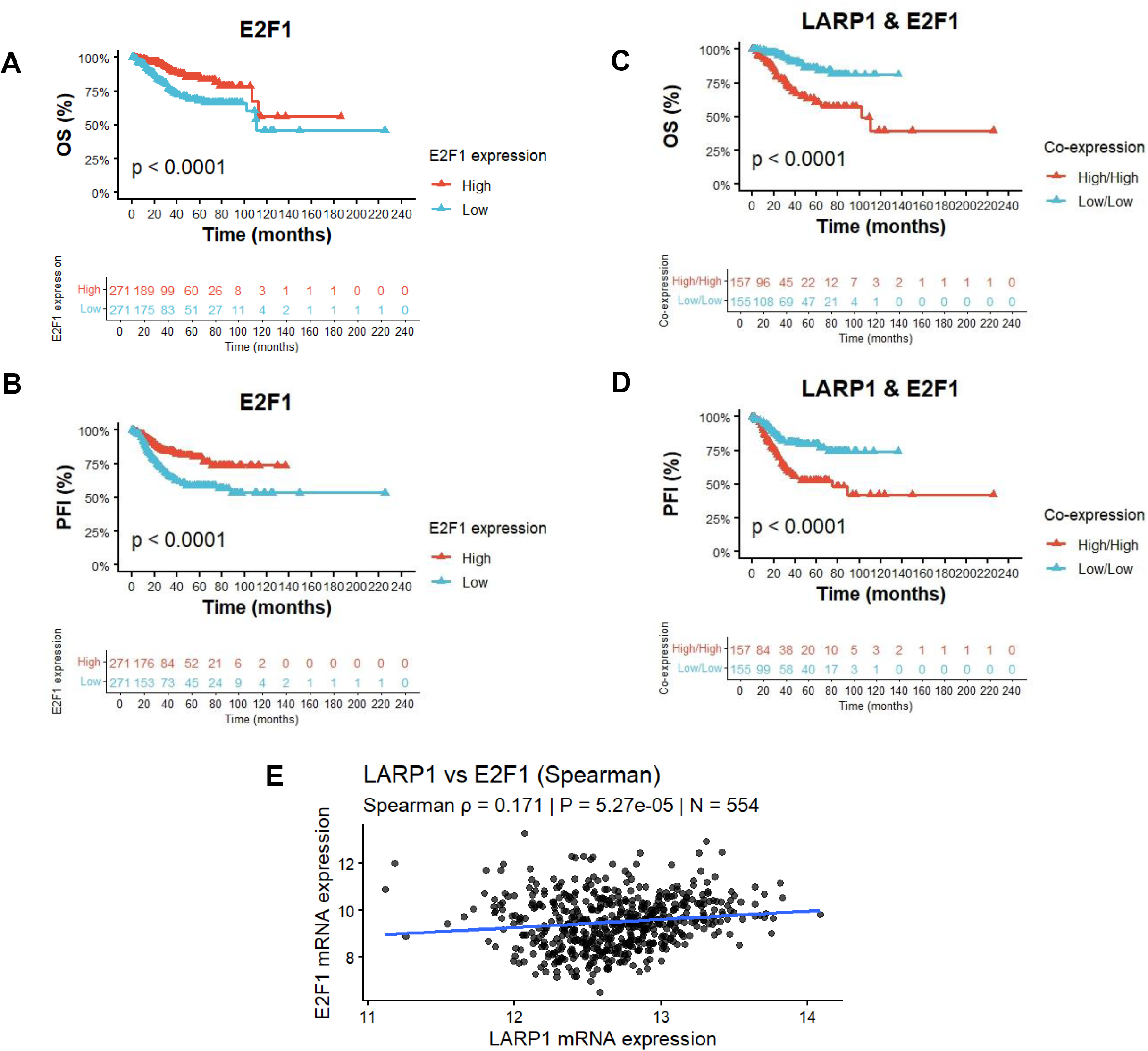
LARP1 positively correlates with E2F1 and co-overexpression of LARP1 and E2F1 is associated with shorter overall survival and progression-free interval in endometrial cancer cohort. (A, B) Kaplan–Meier plot showing that overexpression of E2F1 is associated with shorter overall survival (A) and progression-free interval (B) in endometrial cancer cohort. (C, D) Kaplan–Meier plot showing that co-overexpression of LARP1 and E2F1 is associated with shorter overall survival (C) and progression-free interval (D) in endometrial cancer cohort. (E) Spearman correlation analysis shows a correlation between LARP1 and E2F1 in endometrial cancer patients.

**Figure 4.**
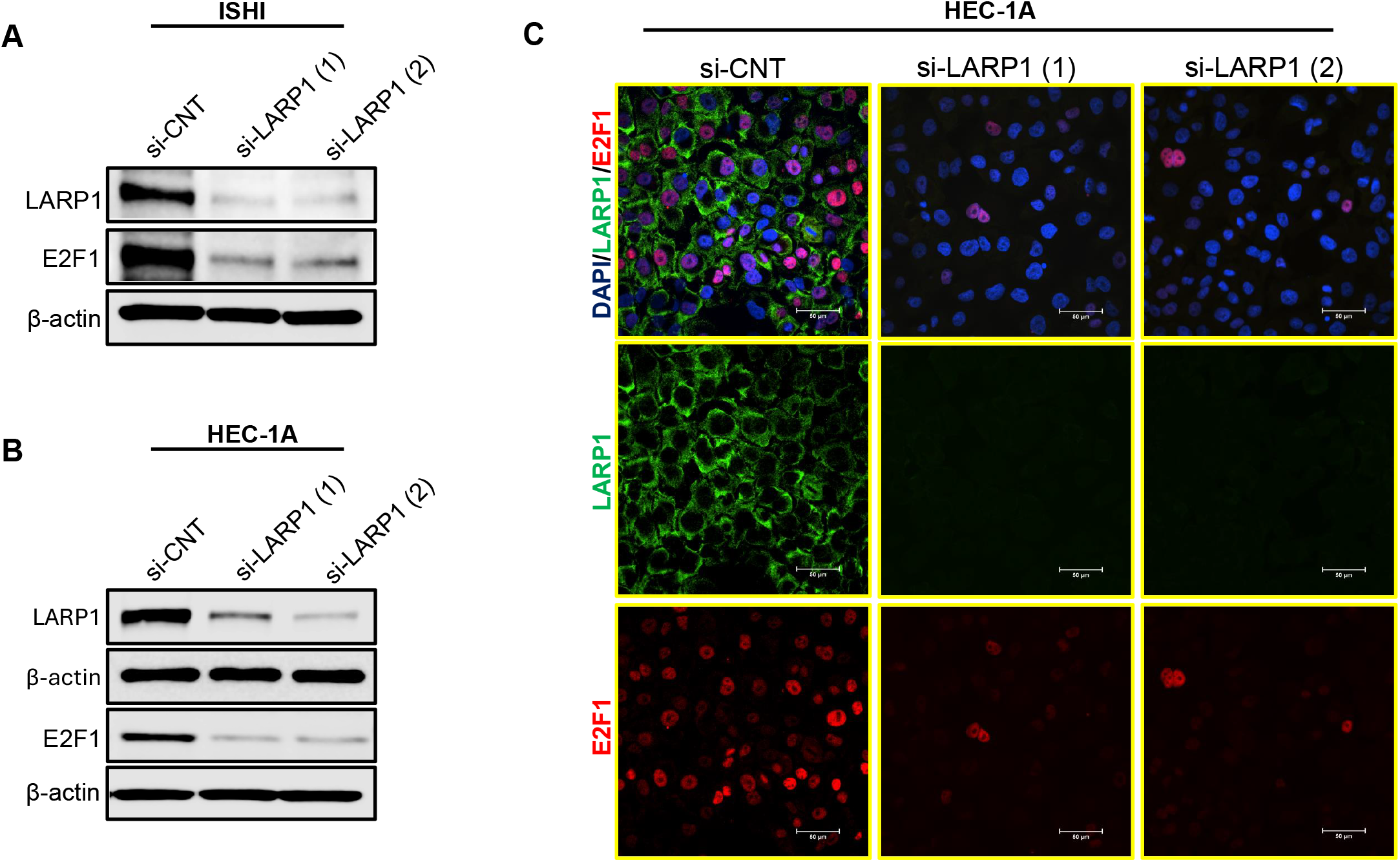
LARP1 knockdown inhibits E2F1 protein expression in endometrial cancer cell lines. (A, B) Immunoblot analysis showing the protein expression of LARP1 and E2F1 after transfecting ISHI (A) and HEC-1A (B) cells with control or LARP1 siRNA. β-actin was used as a loading control. (C) Images representing immunofluorescence staining of LARP1 (green) and E2F1 (red) in HEC-1A cells after transfection with control or LARP1 siRNA. DAPI was used as a counter stain. Scale bar = 50 µm.

### 3.4. LARP1 knockdown sensitizes the response to carboplatin *in vitro*

The combination of carboplatin and paclitaxel represents the standard first-line chemotherapy regimen for the treatment of endometrial cancer patients [1, 3]. We set out to investigate whether LARP1 knockdown sensitizes the response to carboplatin. First, we performed MTS cell viability assay to determine the effect of using escalating doses of carboplatin on the viability of ISHI and HEC-1A. Dose-response analysis revealed that the half-maximal inhibitory concentration (IC_50_) of carboplatin was approximately 48 µM and 97 µM in ISHI and HEC-1A cells, respectively **(Figure 5A, B)**.

**Figure 5.**
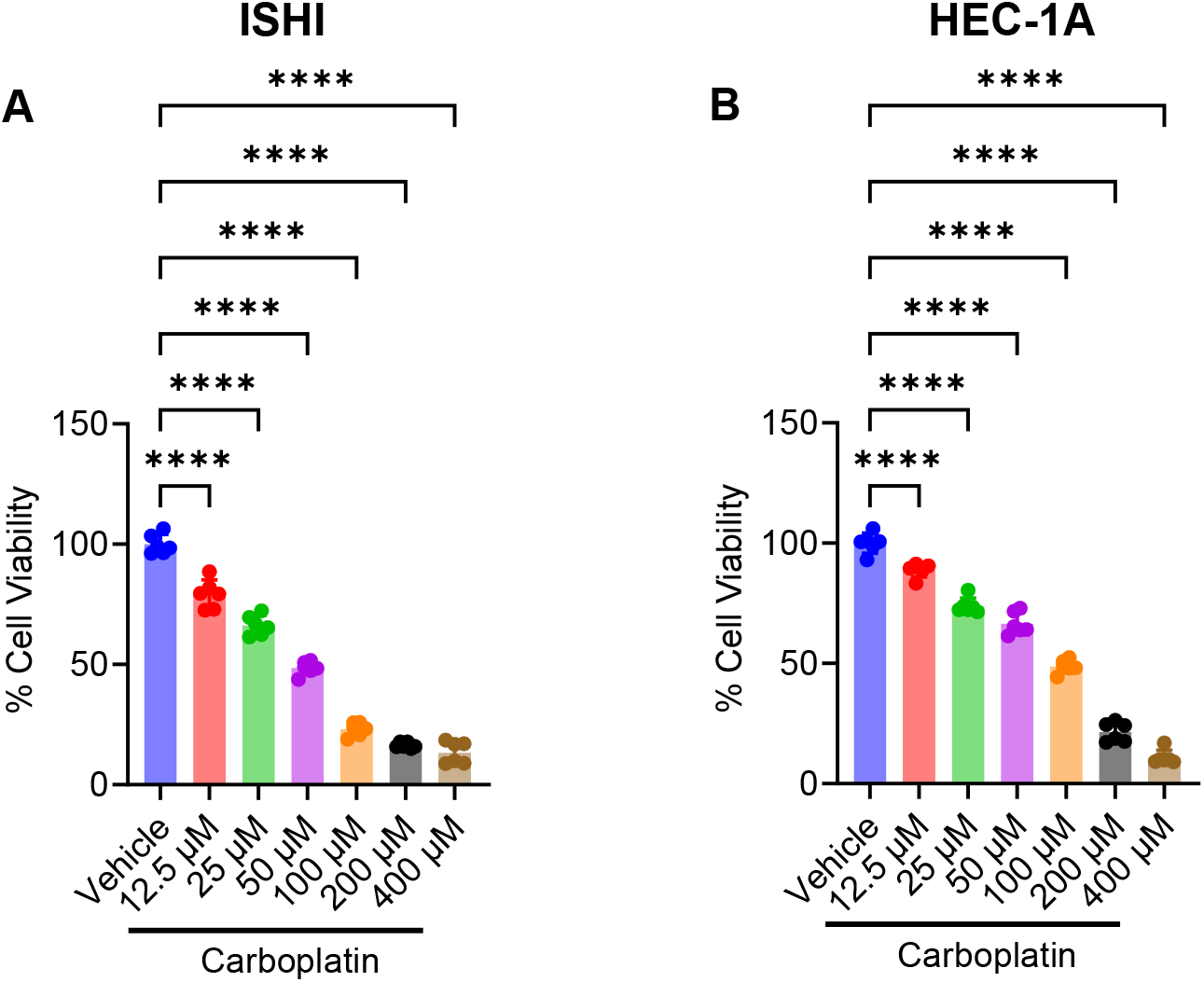
The effect of carboplatin on the viability of ISHI and HEC-1A endometrial cancer cell lines. (A, B) MTS cell viability assay showing the effect of escalating concentration of carboplatin on the viability of ISHI (A) and HEC-1A cell lines. Sterile water was used as a vehicle control. One-way analysis of variance (ANOVA) was used to assess statistical significance among groups, followed by Tukey’s post hoc test for multiple pairwise comparisons. Data are presented as mean ± SD, with individual data points shown as dots.

Based on these findings, we selected a carboplatin concentration (12.5 µM) that exhibited relative resistance in both cell lines and combined it with LARP1 knockdown to assess the impact of such combination on long-term clonogenic survival. Colony formation assay demonstrated that LARP1 knockdown by siRNA 1 and 2 significantly reduced clonogenic survival in ISHI cells, to 50% and 29%, respectively, compared to control siRNA **(Figure 6A, B)**. Similar results were also obtained in HEC-1A cells, where LARP1 knockdown by siRNA 1 or 2 reduced colony formation to 46% and 37% compared to control siRNA **(Figure 6C, D)**.

**Figure 6.**
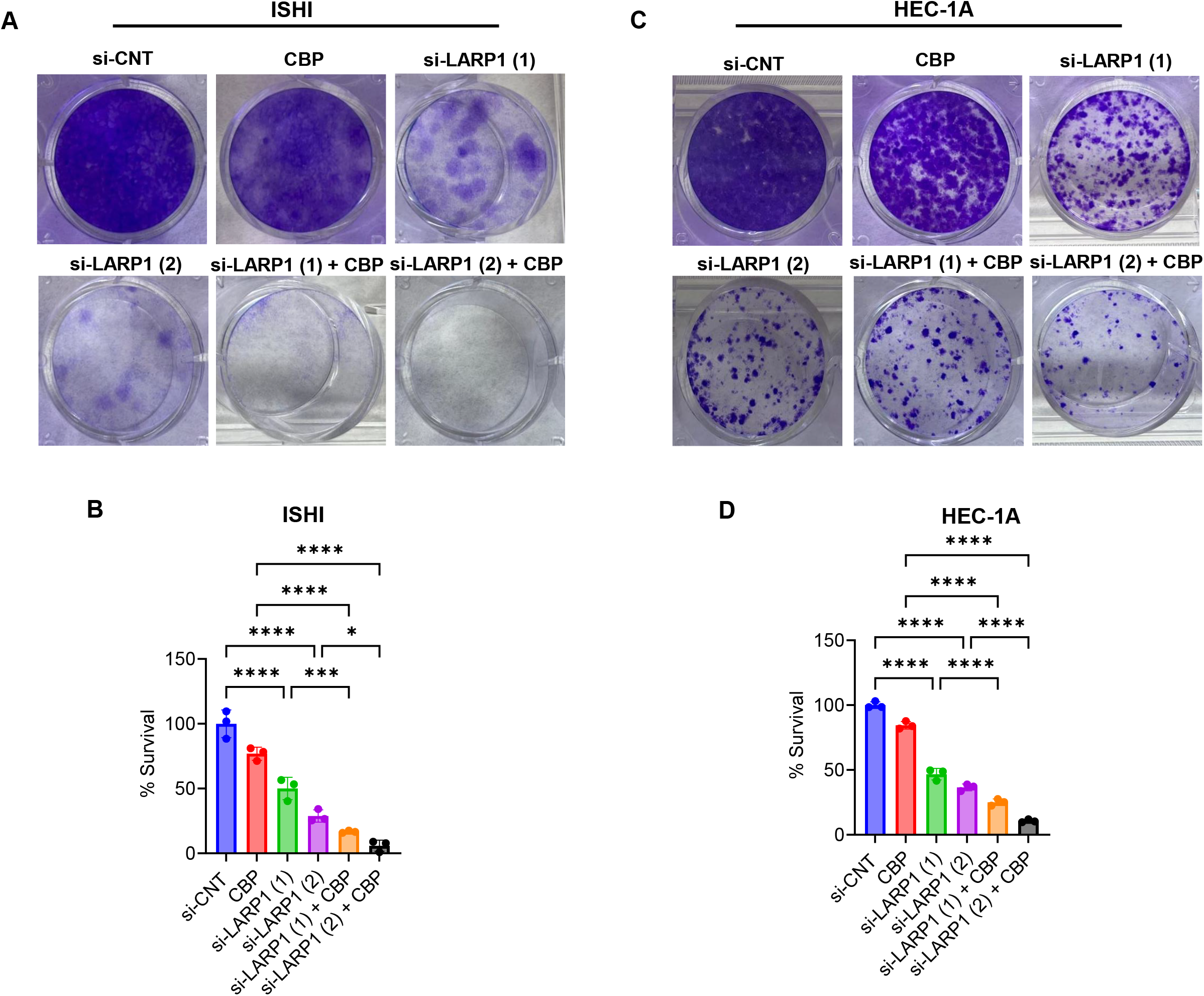
LARP1 knockdown enhances sensitivity to carboplatin *in vitro*. (A, C) Micrographs representing colony formation assay showing the effect of LARP1 transfection and carboplatin treatment on the clonogenic survival of ISHI (A) and HEC-1A (B) cell lines. (B, D) Quantitative analysis of clonogenic assay showing the effect of LARP1 knockdown and carboplatin treatment on the clonogenic survival of ISHI (B) and HEC-1A (D) cells. One-way analysis of variance (ANOVA) was used to assess statistical significance among groups, followed by Tukey’s post hoc test for multiple pairwise comparisons. Data are presented as mean ± SD, with individual data points shown as dots.

Remarkably, combining LARP1 knockdown and carboplatin further suppressed the clonogenic survival in both cell lines. In ISHI cells, the combination of carboplatin and LARP1 siRNA 1 or 2 remarkably decreased colony formation to 17% and 6%, respectively, compared to the corresponding controls **(Figure 6A, B)**. Similarly, in HEC-1A cells the combined treatment regimen carboplatin and LARP1 siRNA 1 or 2 significantly suppressed the clonogenic survival to 25% and 11%, respectively, compared to the corresponding controls **(Figure 6C, D)**. Collectively, these data suggest that LARP1 knockdown could be utilized to enhance the sensitivity to carboplatin-based regimens in endometrial cancer.

## 4. Discussion

In this study, we investigated the functional and clinical significance of LARP1 in endometrial cancer using TCGA patient data and *in vitro* functional studies in HEC-1A and ISHIKAWA cell lines. Our findings suggest that LARP1 promotes tumor cell survival and chemoresistance, and that targeting LARP1 may represent a potential therapeutic strategy to improve response to carboplatin-based chemotherapy in endometrial cancer.

Analysis of the TCGA endometrial cancer cohort revealed that LARP1 overexpression was associated with reduced OS and PFI. Although this association did not remain statistically significant after adjustment for age and stage in multivariate Cox regression analysis, the Kaplan–Meier survival analysis suggests that LARP1 may contribute to aggressive tumor biology. The RNA-binding protein LARP1 has been shown to regulate mRNA stability and translation, particularly transcripts associated with cell growth and proliferation [18, 19]. Several studies have revealed that overexpression of LARP1 is associated with poor prognosis in multiple cancers including ovarian, cervical, and lung cancers [15, 17]. Our findings extend these observations to endometrial cancer and suggest that LARP1 may contribute to tumor progression in this disease.

Furthermore, our findings revealed that LARP1 knockdown significantly decreased cell viability in both ISHI and HEC-1A cell lines, suggesting that LARP1 plays a key role in maintaining endometrial cancer cell proliferation and survival. These findings are in accordance with previous studies showing that LARP1 promotes cancer cell growth through regulation of translation of mRNAs encoding proteins involved in cell cycle progression and metabolic control [17, 19].

Mechanistically, we observed that LARP1 depletion induced apoptosis, as evidenced by increased levels of cleaved PARP1 and cleaved caspase-3. Activation of these apoptotic markers suggests that LARP1 promotes survival signaling in endometrial cancer cells. Previous reports have shown that LARP1 regulates the translation of 5′-terminal oligopyrimidine (5′TOP) mRNAs downstream of mTOR signaling, which encode proteins essential for ribosome biogenesis and cell survival [18, 19]. Furthermore, LARP1 has been shown to interact with the 3′ untranslated regions of *BCL2 and BIK*, stabilizing *BCL2* but destabilizing *BIK* with the net effect of resisting apoptosis [17].

Interestingly, the ISHIKAWA cell line showed higher sensitivity to LARP1 knockdown than HEC-1A cells, as demonstrated by stronger effects on cell viability and apoptosis markers. These differences may be attributed to underlying molecular heterogeneity between endometrial cancer subtypes. ISHIKAWA cells represent a well-differentiated, estrogen receptor–positive endometrioid model, whereas HEC-1A cells display a more aggressive phenotype [20]. Differential dependence on LARP1-regulated translational programs may therefore contribute to variations in therapeutic vulnerability across endometrial cancer subtypes.

In this study, immunofluorescence and western blot analyses showed that LARP1 knockdown inhibited E2F1 protein expression. Furthermore, our TCGA analysis further supports the clinical relevance of E2F1 in endometrial cancer, showing that E2F1 overexpression is associated with shorter OS and PFI, and that E2F1 remains significantly associated with reduced PFI after adjustment for age and stage. Notably, further analysis of TCGA data revealed that co-overexpression of LARP1 and E2F1 is associated with a further reduction in OS and PFI. Additionally, LARP1 mRNA expression levels showed a positive, albeit modest, correlation with E2F1 mRNA levels within the same cohort. E2F1 is a key transcription factor that regulates genes involved in cell cycle progression, DNA replication, and apoptosis [21]. Aberrant regulation of E2F1 has been shown in multiple cancers and is associated with shorter survival rates and poor clinical outcomes [22, 23]. Taken together, these data suggest that LARP1 and E2F1 may function cooperatively to enhance endometrial cancer progression. The precise mechanism underlying the crosstalk between LARP1 and E2F1 remains to be elucidated. However, it is plausible that LARP1 enhances the stability or translation of E2F1 transcript, given the well-known function of LARP1 as an RNA-binding protein [17, 24]. Alternatively, LARP1 may regulate the translation of upstream regulators of cell cycle to indirectly regulate E2F1 expression [14]. Further studies are warranted to precisely determine the exact mechanism by which LARP1 regulates E2F1.

An important clinically relevant finding of the current study is that LARP1 knockdown enhances sensitivity to carboplatin treatment. Carboplatin is the cornerstone for the management of advanced or recurrent endometrial cancer; however, treatment resistance is a major clinical limitation [5]. Overexpression of E2F1 has been shown to enhance platinum resistance by upregulating DNA damage response and inhibiting apoptosis [25]. We speculate that LARP1 may contribute to chemotherapy resistance and that targeting LARP1 could induce apoptosis and inhibit E2F1 expression, which in turn may enhance the sensitivity to platinum-based therapy.

This study has multiple limitations. First, we conducted the clinical association analyses using retrospective TCGA datasets, so we need to validate the prognostic value of LARP1 in independent clinical groups. Second, while our lab experiments showed that LARP1 plays a role in endometrial cancer cell survival and chemotherapy response, establishing *in vivo* models is needed to validate the beneficial effects of LARP1 targeting. Finally, the molecular mechanisms underlying the regulation between LARP1 and E2F1 are not fully understood and warrant further investigation.

In conclusion, our findings demonstrate that LARP1 promotes tumor survival, inhibits apoptosis, and contributes to resistance to platinum-based chemotherapy. Furthermore, targeting LARP1 could be used as a potential therapeutic approach to mitigate tumor growth and enhance sensitivity to chemotherapy in endometrial cancer.

## 5. Data accessibility

Normalized RNA-seq expression counts (FPKM-UQ) and clinical data for endometrial cancer patients were obtained from The Cancer Genome Atlas dataset via Genomic Data commons Portal. All other data that support the findings of this study are available upon request from the corresponding author.

## 6. Author Contribution

AME conceived the project. AME performed experiments, TCGA analysis, and statistical analysis. AME and MWE wrote the article and assembled the figures. SAS revised the manuscript. All authors reviewed the final manuscript.

## 7. Use of AI tools

Artificial intelligence tools were used to check grammar and improve the flow of the manuscript. AI was not used to generate scientific content, provide references, write the discussion, or determine authorship.

